# Ultra-Content Screening (UCS): Toward the Big Blood Picture

**DOI:** 10.64898/2026.01.20.700546

**Authors:** Sebastian Bernhard Kloubert, Simon Renders, Oliver Bannach, Thomas Dino Rockel, Andreas Trumpp, Dieter Willbold

**Affiliations:** Institut für Physikalische Biologie (IPB), Heinrich-Heine-Universität Düsseldorf, Düsseldorf, Germany; Institute of Biological Information Processing, Structural Biochemistry (IBI-7), Forschungszentrum Jülich, Jülich, Germany; Heidelberg Institute for Stem Cell Technology and Experimental Medicine (HI-STEM gGmbH), Heidelberg, Germany; Division of Stem Cells and Cancer, German Cancer Research Center (DKFZ) and DKFZ-ZMBH Alliance, Heidelberg, Germany; German Cancer Consortium (DKTK), German Cancer Research Center (DKFZ), Heidelberg, Germany

**Keywords:** Single-cell proteomics, immunocytochemistry, cyclic analysis, iterative staining, cell based diagnostics

## Abstract

We present Ultra-Content Screening (UCS), a novel, scalable method combining cyclic immunostaining with high-dimensional image-based single-cell proteomics.

UCS utilizes fluorescein isothiocyanate-conjugated antibodies and iterative staining-photobleaching cycles to analyze up to 40 markers in up to 100,000 peripheral blood mononuclear cells per experiment. Through precise image registration, nuclear segmentation, signal harmonization, and normalization, UCS ensures the robust tracking of individual cells across all staining cycles. Data analysis via SPADE trees allows qualitative evaluation of expression patterns and cellular phenotypes. Application to samples from acute myeloid leukemia patients demonstrates UCS’s potential to uncover disease-specific expression profiles and immune subpopulations. The method provides unprecedented depth in single-cell proteomic analysis of blood samples, offering valuable insights for diagnostics, personalized medicine, and therapeutic approaches.

## 1. Introduction

The combination of immunocytochemistry and big data analysis has the potential to reveal previously concealed information. Traditional immunostaining methodologies allow the identification of mean protein expression profiles across cell populations. This approach obscures individual cell characteristics and essential information about cellular heterogeneity. By integrating high-dimensional single-cell analysis with big data analysis methods, it is possible to characterize the combined molecular properties of individual cells across entire populations to identify rare cell subpopulations and subtle expression patterns that would remain undetected in population-based approaches. In recent years, new technologies have been developed in the field of single-cell proteomics that attempt to represent the heterogeneity of cells, with different approaches and varying depths of information. [1] [2] [3] In the current landscape of cellular phenotyping, the majority of established methods are tissue-based. [4][5][6] When cyclic staining protocols are applied to achieve higher multiplexing, they typically require the use of costly reagents—such as DNA-conjugated antibodies or specialized chemical bleaching agents—or necessitate access to sophisticated instrumentation, including mass spectrometers or custom-designed imaging platforms. [5][6][7] [8] These constraints present relevant barriers to the routine adoption of highly multiplexed single-cell analyses in standard diagnostic workflows.

Our Ultra-Content Screening (UCS) method integrates the concept of cyclic immunostaining with the photosensitive fluorophore fluorescein isothiocyanate (FITC) and subsequent photobleaching and immobilization of immune cells on glass carriers and allows the acquisition and analysis of data from 40 antibody stains for up to 100,000 peripheral blood mononuclear cells (PBMC) in a single experiment. [9] Basic laboratory equipment and an LED microscope are all that are needed for the acquisition of stained images and the bleaching of the signal directly afterwards in the same device. Stacking the acquired staining images allows the application of modern big data analysis methodologies, for example, qualitative evaluation by using Spanning-Tree Progression Analysis of Density-normalized Events (SPADE) trees. (Figure 1) [10] The depth of information obtained through this method is unparalleled by any currently known technique on this scale. The analysis of this combined molecular information offers the potential to facilitate a more detailed characterization of disease markers. The ability to detect disease-specific cellular signatures in potential presymptomatic stages provide novel perspectives for preventive intervention strategies and personalized treatment concepts. This potential can be utilized to improve lineage determination and subclassification in diseases such as acute myeloid leukemia (AML), as well to provide more information about the differentiation and distribution of leukemia cells. AML is an aggressive malignant disease of the blood system characterized by the clonal expansion of immature myeloid cells that are unable to differentiate properly, leading to impaired hematopoiesis. [11] Although morphology and cytogenetics remain crucial, multiparametric flow cytometry is essential for the diagnosis of AML, as it allows detection of specific immunophenotypic patterns in leukemic blasts. [12] The implementation of our new ultra-content screening method has the potential to enhance this process. In this article, we illustrate the implementation of the method and present example results derived from data sets originating from 11 AML patient samples.

**Figure 1:**
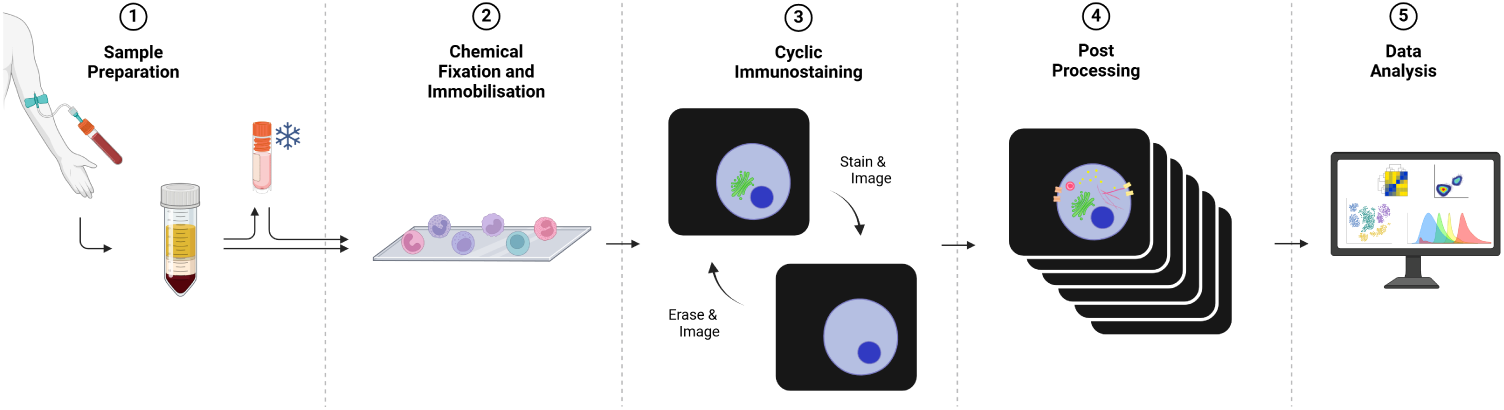
Schematic workflow of the Ultra-Content Screening Method. (1) Blood samples are collected and mononuclear cells are isolated via density gradient centrifugation. (2) Blood cells are chemically fixed and immobilized on glass plates. (3) Iterative immunostaining via antibody staining, signal capture and signal elimination by photobleaching. (4) Image data are processed by registration, segmentation, harmonization and normalization. (5) Extended data analysis including qualitative evaluation of the results via SPADE tree visualization.

## 2. Methods

### 2.1. Sample Preparation

This protocol can be used to prepare whole blood samples as well as buffy coats to isolate the mononuclear fraction of leukocytes (peripheral blood mononuclear cells, PBMCs) and freeze for medium-term storage at −80 °C or long-term storage in liquid nitrogen or transport. The use of buffy coats results in a higher yield. The complete list of chemicals and equipment necessary for the protocol can be found in Table 1. These steps take approximately 90 minutes, and there is no opportunity to pause.

1. Collect PB/BM samples from heparin vacutainer (c. 7.5 mL) into 50 mL Falcon tube with a 100 µm filter using stripette
2. Wash collection tubes twice with PBS and fill Falcon up to 35 mL with PBS
3. Add 15 mL Ficoll-Paque to new Falcon tubes
4. Pour slowly BM solution on top of the Ficoll-Paque solution with max 35 mL of sample solution per Falcon (sample 1:2 to Ficoll-Paque)
5. Centrifuge at 400 g x 30 min at RT, with deactivated breaks **Note:** Make sure centrifuge is at RT before starting the centrifugation or cell separation may be affected
6. Aspirate 15 mL from the top of the 50 mL Falcon tube to remove plasma and platelets
7. Collect the buffy coat to a new 50 mL Falcon tube, and then the ficoll up to the RBC level
8. Fill the collection tubes with PBS
9. Centrifuge at 500 g x 8 min and resuspend in 2% FCS/PBS
10. Pool vials together and count the cells
11. Incubate cells in FCS for 5 min at RT and then for 10 min at 4 °C
12. Add appropriate volume of FCS/20% DMSO to make final DMSO concentration in freezing media 10% (e.g. 5 mL FCS + 5 mL FCS/20% DMSO)
13. Prepare aliquots of 5 - 25 × 10^6^ cells in 1 mL and place cryotubes in Mr. Frosty **Note:** Make sure to have several tubes per patient; rather more vials than more cells per vial
14. Transfer Mr. Frosty to −80 °C
15. Transfer freezing tubes after 1 day into liquid nitrogen tanks for longer storage

**Table 1:** Chemicals and equipment needed for the sample preparation protocol.

| Chemicals | Equipment |
| --- | --- |
| • Histopaque<br>• PBS<br>• PBS + 2% FCS<br>• FCS<br>• FCS + 20% DMSO | • Centrifuge<br>• Cryotubes<br>• Freezing Container |

### 2.2. Thawing

Thaw samples in preparation for chemical fixation. The PBS + FCS solution in Step 3 should be slowly added to slow the concentration changes and avoid clumping. These steps take approximately 30 minutes and there is no opportunity to pause. The subsequent fixation protocol (see Section 2.3) should be performed immediately afterward. The complete list of chemicals and equipment necessary for the protocol can be found in Table 2.

1. Prepare by cooling the centrifuge to 4 °C and heating 15 mL of PBS + 10% FCS in a 37 °C water bath
2. Thaw cryo vials in a 37 °C water bath for 90 s or until the sample is thawed 70%
3. Transfer sample to PBS + 10% FCS
4. Centrifuge at 300 g for 5 min at 4 °C
5. Discard supernatant and resuspend in 3 mL PBS + DNase I (100 µg/mL)
6. Incubate at RT for 15 min
7. Add 12 mL PBS + 10% FCS

**Table 2:**
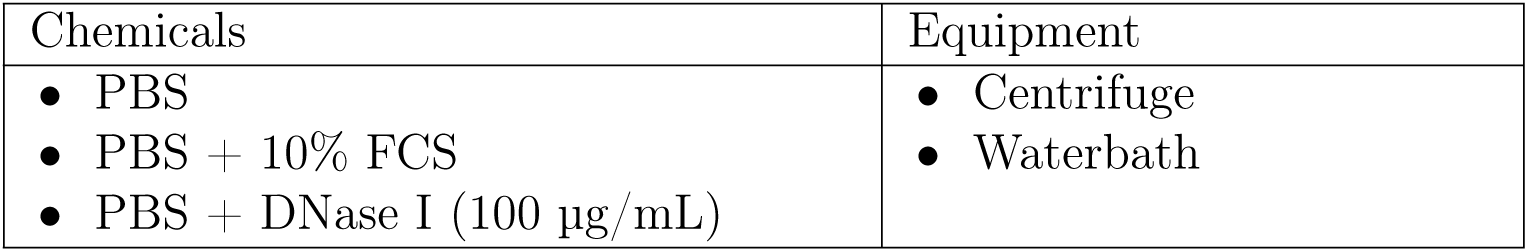
Chemicals and equipment needed for the thawing protocol.

### 2.3. Fixation

To prevent the biological properties of the cells under study from changing over the time of the cyclic immunostaining, the sample is chemically fixed. In addition, the nuclei are stained with Hoechst 33342. These steps take approximately one hour. The experiment may be interrupted from the first wash step in step 5, but should be completed as soon as possible. After staining the nuclei in step 7, the sample should be incubated in the dark. After completion, samples can be stored at 4 °C for an intermediate period of time. The complete list of chemicals and equipment necessary for the protocol can be found in Table 3.

1. Centrifuge 300 g, 5 min @ RT
2. Discard supernatant and resuspend in 10 mL PBS + 4% PFA
3. Incubate 15 min @ RT
4. Centrifuge 300 g, 7 min @ RT
5. Discard supernatant and resuspend in 50 mL PBS
6. Centrifuge 300 g, 7 min @ RT
7. Discard supernatant and resuspend in 10 mL PBS + Hoechst 33342 (10 µg/mL)
8. Incubate 30 min @ RT, in the dark
9. Centrifuge 300 g, 7 min @ RT
10. Discard supernatant and resuspend in 50 mL PBS
11. Repeat Steps 9 & 10
12. Count cells

**Table 3:** Chemicals and equipment needed for the fixation protocol.

| Chemicals | Equipment |
| --- | --- |
| • PBS<br>• PBS + 4% PFA<br>• PBS + Hoechst 33342(10 µg/mL) | • Centrifuge |

### 2.4. Immobilization

In order to assign the information from immunostaining of the different cycles to the same cell, the cells will be immobilized on imaging plates. In Step 3, the solution must be applied with precision, ensuring that the drop is precisely positioned in the center of the well and that the liquid does not come into contact with the edges. It is crucial to note that the working volume of the plate, with the entirety of the bottom covered, is merely 300 µL. This is only marginally greater than the 200 µL utilized. It is crucial to avoid liquid entering the edges, a significant proportion of the cells will become immobilized at the well edge, resulting in increased scattering and reduced accessibility for subsequent analysis. The immobilization process is approximately one hour and thirty minutes in duration and should not be interrupted during this time. Subsequent to this, the wells should be washed five times with 700 µL of phosphate-buffered saline (PBS) each, in order to evaluate which well has immobilized the most cells and the initial cell recovery. Afterwards the plate can be stored at 4°C covered with aluminum foil. Immunostaining should be carried out over the next few days. The complete list of chemicals and equipment necessary for the protocol can be found in Table 4.

1. Centrifuge sample volume equivalent to 8 × 10^6^ cells 300 g, 7 min @ RT
2. Discard supernatant and resuspend in 1.6 mL PBS
3. Add 200 µL in each of the eight inner wells of MACSwell 24 Imaging Plates
4. Incubate for 1 hour @ RT, in the dark
5. Centrifuge 10 min @ 200 rpm
6. Add 700 µL PBS, aspirate completely and add 300 µL PBS + 4% PFA
7. Incubate 15 min @ RT, in the dark
8. Aspirate and add 1 mL PBS

**Table 4:** Chemicals and equipment needed for the immobilization protocol.

| Chemicals | Equipment |
| --- | --- |
| • PBS<br>• PBS + 4% PFA | • Centrifuge<br>• LED Microscope<br>• Imaging Plates |

### 2.5. Immunostaining

The final and most tedious phase of the laboratory section is the cyclic staining of the samples. The photosensitivity of the fluorophore fluorescein isothiocyanate (FITC) enables the removal of the antibody staining signal after its acquisition in the microscope by photobleaching, thereby allowing the initiation of the staining process anew. In our experimental setup, 40 staining cycles were executed for each data set. The total duration of one cycle, including the 30-minute staining incubation, the 15-minute bleaching phase, and the subsequent washing and handling of the plate, is between 50 and 55 minutes. It is important to note that, in principle, the process can be paused after each cycle and resumed the subsequent day. Ideally, all dyeing cycles should be completed within one week and not interrupted by a weekend to avoid cell loss. The complete list of chemicals and equipment necessary for the protocol can be found in Table 5.

1. Completely remove buffer and add 300 µL PBS + 2% antibody.
2. Incubate 30 min @ RT, in the dark
3. Add 700 µL PBS
4. Wash 4 times with 700 µL PBS and incubate for 30 s in between
5. Add PBS to a volume of 1 mL
6. Perform first measurement with LED microscope followed by 15 min of photobleaching
7. Perform second measurement in LED microscope

**Table 5:** Chemicals and equipment needed for the immunostaining protocol.

| Chemicals | Equipment |
| --- | --- |
| • PBS<br>• Antibody recombinant human IgG1 fluorescein isothiocyanate (FITC) | • LED Microscope |

### 2.6. Postprocessing

#### Registration

The insertion of the plate into the microscope on multiple occasions results in a minor, unavoidable variation in the plate’s position for each measurement. In order to directly compare the images of different cycles on pixel basis, this variance must be corrected. To this end, a cross-correlation technique is employed, relying on a subpixel registration algorithm. [13] The registration of images between cycles and within a cycle, prior to and following bleaching, ensures optimal comparability.

#### Segmentation

After ensuring that all images are correctly aligned, it is essential to acquire data regarding cell position to perform the analysis. For this purpose, the cells are spatially segmented based on the measured signal of the Hoechst 33342 nuclear stain. Initially, the potential excess signal from uneven illumination is reduced through implementation of a rolling ball algorithm. [14] Subsequently, Voronoi-Otsu labeling is performed. [15] For this purpose, the results of two separate steps are combined. In the first step, a Gaussian blur is applied to the image, and then the image is reduced to its maximum points. [16] In the second step, another Gaussian blur is applied and an Otsu thresholding is applied to generate a binary image. [16][17] The combination of the binary image with the maximum points and the application of a Voronoi diagram results in a segmentation of cell nuclei in which each cell is assigned a distinct identity.[18][19] This approach ensures that individual cells can be specifically addressed during subsequent analysis and that the signal intensity of antibody stainings can be mapped across different cycles.

#### Harmonization

Cells are considered for analysis only if they are found at the same coordinates in all staining cycles. To this end, the center of mass of each cell is calculated, and for each cycle, it is verified that the center of mass is within the segmentation mask of the respective nuclear stain. In the event that the center of mass does not overlap with the segmentation mask, the cell is excluded from the overall analysis.

#### Normalization

The photobleaching of the antibody fluorophore results in residues that should be excluded from subsequent cycle analysis. Therefore, the measurement result of each cycle is normalized by subtracting the residual signal from the previous bleach. This ensures that only the signal that was added by subsequent antibody staining is included in the analysis.

### 2.7. Analysis

Given that the analysis is now constrained to consider only cells that have undergone successful immobilization in all cycles and that only the signal of the last antibody staining is considered, the signal intensity of each cell can now be measured. The median value of the segmented area of the cell is utilized as the signal intensity per cell and staining cycle. The utilization of rudimentary graphs proves insufficient to represent the complexity of the resulting data sets. The Spanning-Tree Progression Analysis of Density-normalized Events (SPADE) is a method that allows for the qualitative evaluation of the results. This method was developed for the purpose of visualizing a large number of multiparametric single-cell data sets. It combines density-dependent downsampling, hierarchical clustering, and minimum spanning trees (MST). By reducing the data complexity and subsequent clustering, SPADE enables a structured visualization of the cell hierarchy and supports the identification of biological phenotypes within the data set. [10]

## 3. Results

Although 1 × 10^6^ cells were applied per well, the number of cells that were actually immobilized on the plate varied considerably. Box plot analysis of the data showed that the number of immobilized cells ranged from a minimum of 21,797 to a maximum of 101,123. The median was 56,303 cells, while the first quartile was 41,821 cells and the third quartile was 79,728 cells. (Figure 2 a)

**Figure 2:**
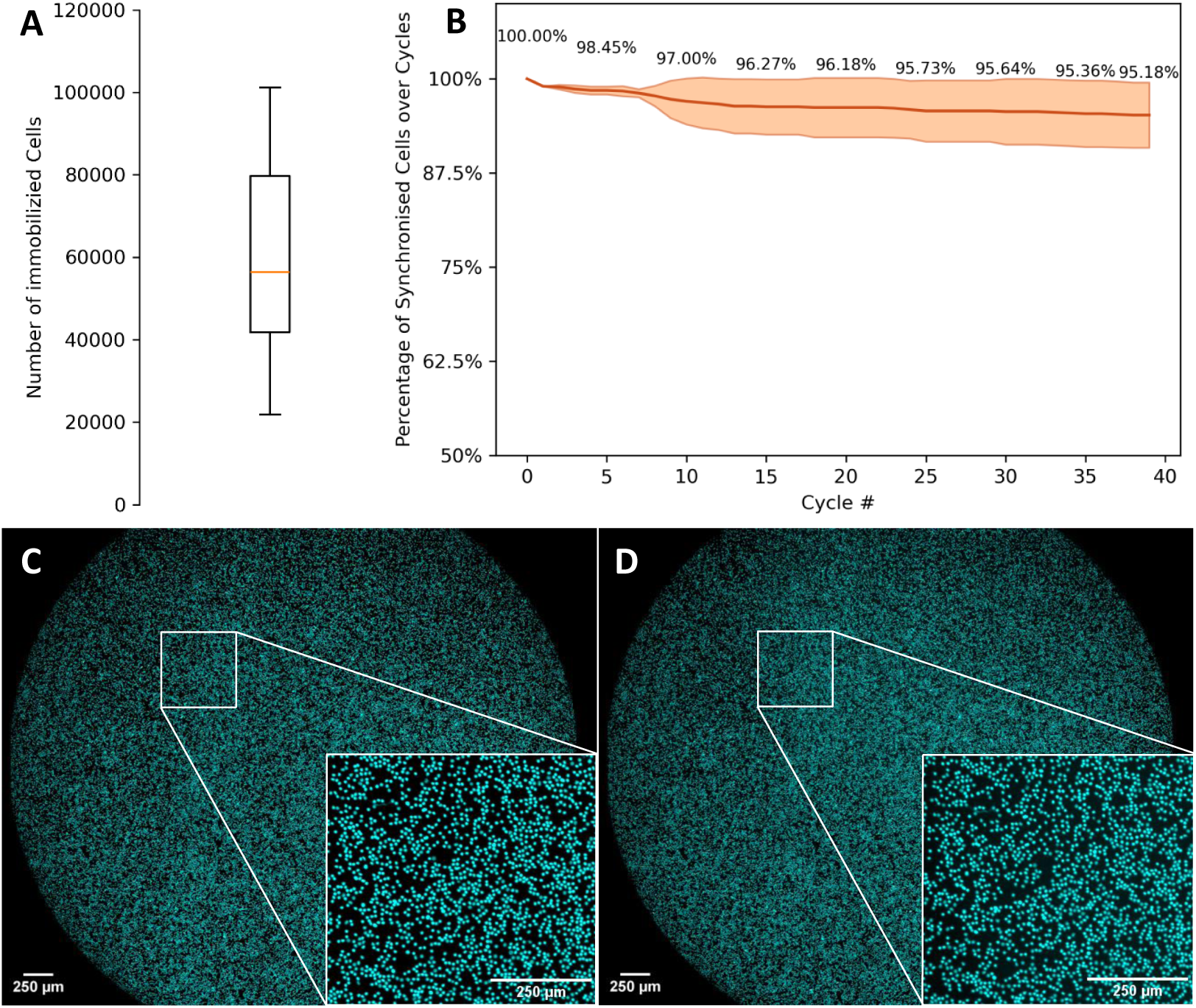
Cell immobilization and recovery over 40 cycles. (a) Boxplot of initial immobilized cells after five washing steps of 11 patient samples. (b) Percentage of remaining cells over the course of 40 staining cycles in 11 patient samples. (c-d) Comparison of the microscope fields-of-view of the core stain after 1 and 40 staining cycles.

After immobilizing the cells on the plate and subsequent initial washing steps, a stable cell count was observed throughout all staining cycles. No significant losses of immobilized cells occurred, so that the initial cell count was essentially maintained. Over the course of 40 cycles, an average of 95.18% ± 4.33% of the originally immobilized cells were retained and could be harmonized at the same location across all cycles. The error rate up to the eighth staining was only 0.54%. From the ninth staining onwards, the error rate increased, reaching a value of 3.41% in the twelfth staining and it remained stable afterwards. (Figure 2 b) The signal intensity of the Hoechst 33342 core stainings proved to be largely stable during the experiments. Only slight bleaching was observed, but this did not impair the amount of cells detected by the segmentation algorithm. (Figure 2 c-d, Supplemental Figure **??**)

The cells were successfully labeled with fluorescein isothiocyanate (FITC). (Figure 3) In a representative image sample of 37 cells, 19 cells showed a strong CD3 signal, while a weak CD3 signal was detected in 6 cells. (Figure 3 a-c) With regard to CD4 expression, 13 cells showed a strong CD4 signal and 6 cells showed a weak CD4 signal. (Figure 3 g-i) Analysis of CD8 expression revealed that 9 cells showed a strong CD8 signal and 8 cells showed a weak CD8 signal. (Figure 3 m-o) It was noticeable that cells with strong CD4 or strong CD8 expression were generally mutually exclusive; only in one case was coexpression of strong CD4 and CD8 signals observed. The planned bleaching effect occurred as expected. (Figure 3 e, k) However, the cell with CD4 / CD8 coexpression showed a certain residual fluorescence signal after both bleaching procedures. (Figure 3) A robust algorithm was used to register the image data, ensuring high precision. The offset determined between the images was mostly less than 10 pixels, indicating very good agreement between the image data. The image data was always aligned relative to the first core stain of the first cycle. For safety reasons, registration was also performed between the respective stain and bleach images. However, the deviations observed were negligible. Even in areas with densely packed cells, each cell was reliably assigned its own unique identity. (Supplemental Figure **??**) The good segmentation quality enabled reliable verification of cell positions across the various cycles. It was found that the vast majority of cells remained in the same positions in subsequent cycles. On average, 95.18% of cells were synchronously identifiable across all cycles. (Figure 2 b) After harmonization, the median signal intensities of the synchronous cells were recorded for all 40 antibody stains. The cells were clustered based on similar signal intensities to identify functionally and phenotypically related groups of cells. A minimal spanning tree (SPADE tree) was created to visualize the relationships between the clusters. In this representation, neighboring clusters have similar properties, whereas those that are farther apart have greater differences in their signal profiles. The size of a cluster represents the number of cells with similar properties that are represented by that node.

**Figure 3:**
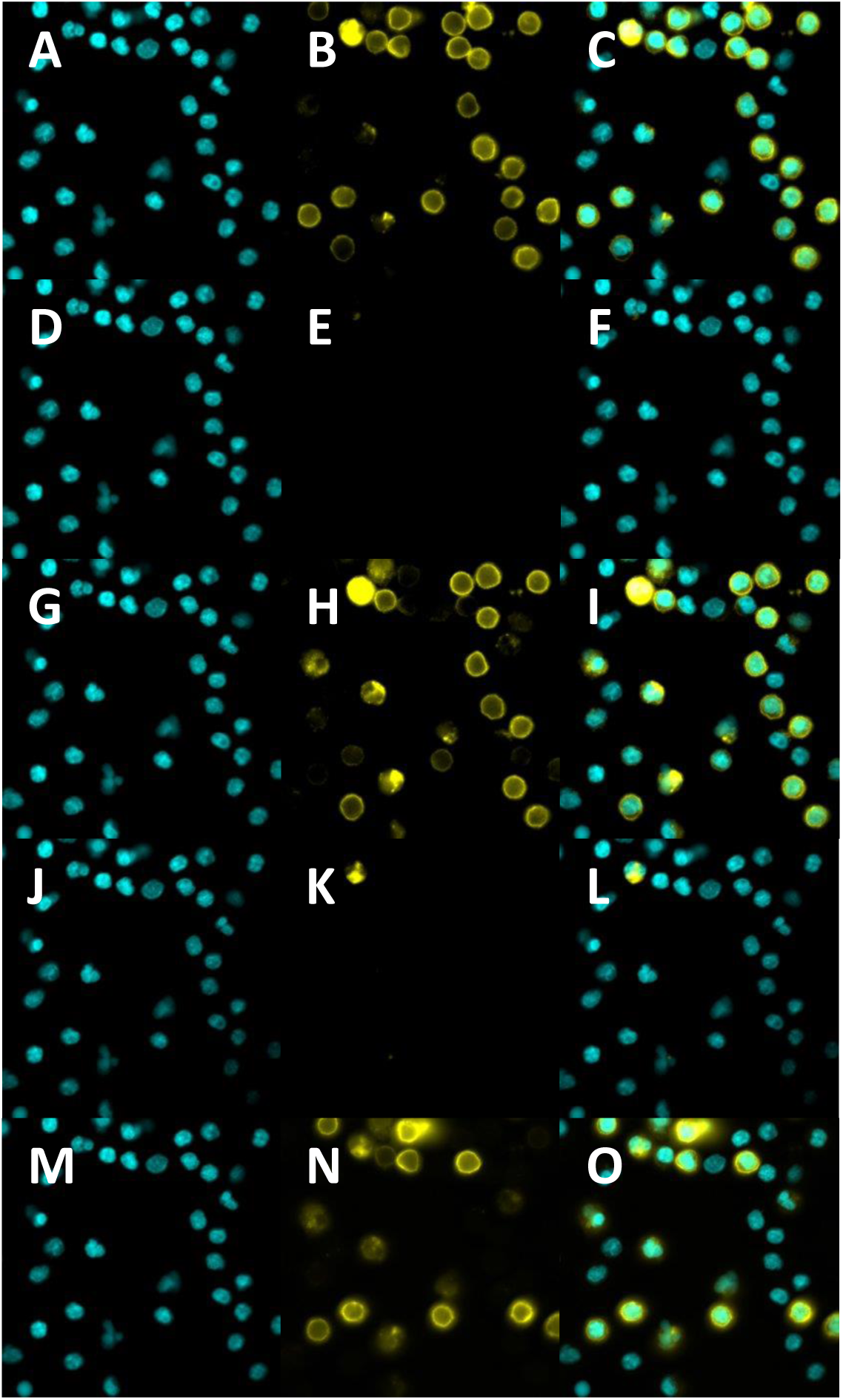
Demonstration of cyclic staining and bleaching. (a-c) Hoechst 33342 nucleus stain signal, FITC antibody stain signal and overlay of CD3 staining. (d-f) Hoechst 33342 nucleus stain signal, FITC antibody stain signal and overlay after 15 minutes of photobleaching. (g-i) Hoechst 33342 nucleus stain signal, FITC antibody stain signal and overlay of CD4 staining. (j-l) Hoechst 33342 nucleus stain signal, FITC antibody stain signal and overlay after 15 minutes of photobleaching. (m-o) Hoechst 33342 nucleus stain signal, FITC antibody stain signal and overlay of CD8 staining. All images show the same field of view and were acquired at 40x magnification.

Larger clusters are located in the center of the SPADE tree, whereas several smaller clusters are located at the edge. (Figure 4 l) Overall, CD3 displays a consistently elevated signal, with only a few central nodes showing low signal intensity. (Figure 4 a) CD4 demonstrates notably elevated signal intensities, particularly on the right side of the SPADE tree, with individual strongly positive nodes also observed on the left. (Figure 4 b) For CD8, medium signal intensities are detected in the upper right area, with an elevated signal at the lowest branch of the tree. (Figure 4 c) CD16b demonstrates a markedly elevated signal in the upper right quadrant, while no signal is detectable in the left portion of the SPADE tree. (Figure 4 d) CD19 protein expression is predominantly observed in the right section of the tree, while moderate signal intensities are detected in the middle. (Figure 4 e) CD33 demonstrates a high signal in the right section of the SPADE tree; in the left section, expression is largely undetectable, with the exception of one branch that shows moderate intensity. (Figure 4 f) The expression of CD56 is high on the right side of the tree, while a heterogeneous, partly high signal distribution can be observed on the left side. (Figure 4 h) CD66b demonstrates notably elevated signal intensities on the right side of the tree and an absence of signal on the left side. (Figure 4 i) For CD123, a very high signal is detected on the upper right side of the SPADE tree, whereas almost no signal is present on the left side. (Figure 4 j) CD163 demonstrates moderate signal intensity in the right area and isolated moderate signals on the lower part of the tree. (Figure 4 k) CD45 expression is consistently high across all clusters. (Figure 4 g)

**Figure 4:**
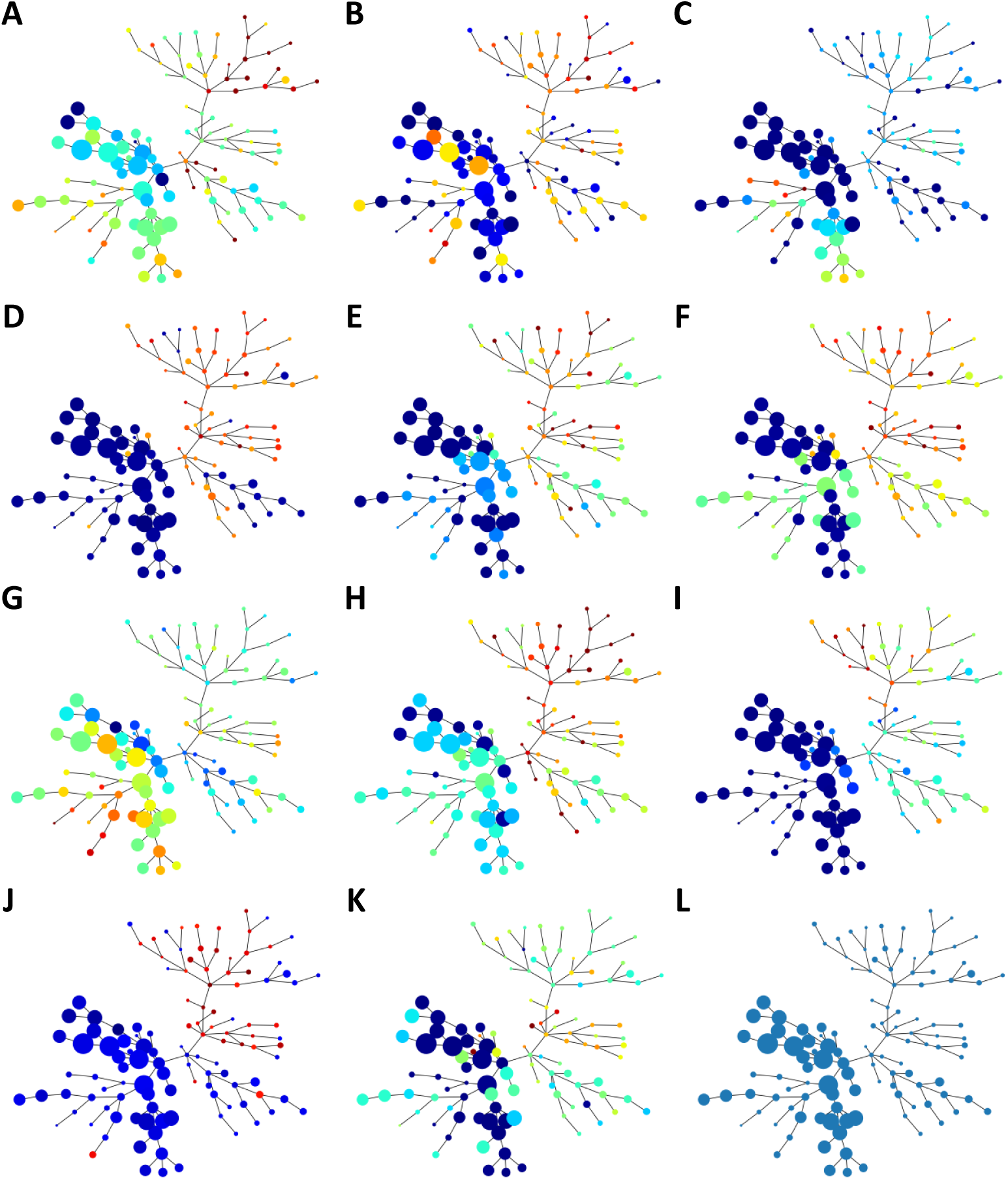
Exemplary qualitative evaluation of the calculated fluorescence data via SPADE trees. Each SPADE tree is a minimal spanning tree, wherein clusters of cells with analogous fluorescence data distributions are proximate to each other. The clusters are color-coded according to the intensity of the mean fluorescence intensity that was measured, ranging from blue for low to red for high for the corresponding antibody staining (a) CD3, (b) CD4, (c) CD8, (d) CD16b, (e) CD19, (f) CD33, (g) CD45, (h) CD56, (i) CD66b, (j) CD123, (k) CD163. (l) Node size represents the number of cells assigned to the respective cluster.

## 4. Discussion

Due to the physical distance between the locations where the samples were taken and where the analysis took place, it was necessary to freeze the cells and thaw them again for further processing. The yield was found to vary notably following thawing; however, a clear cause could not be identified. (Supplemental Table **??**) One approach to minimize these losses would be to directly chemically fix the peripheral mononuclear cells (PBMCs) after they have been isolated from fresh blood or buffy coat, thereby avoiding the freezing and thawing process. However, as long as a minimum of 8 × 10^6^ cells were available per approach, the observed losses do not serve as a limiting factor for the application of the method. The variability in the number of cells initially immobilized on the plate represents a potential factor influencing the comparability between different data sets and should be further analyzed in future studies to improve standardization and reproducibility of the experiments. However, cell recovery is excellent in the further course of the experiment, so that the initial fluctuations in number of cells do not have a significant negative effect on the quality or significance of the subsequent analyses. The segmentation of cell nuclei is not significantly affected by the bleaching process, so that individual cells can still be reliably identified and distinguished. However, to prevent potential loss of contrast and maintain consistently high segmentation quality even after multiple staining and bleaching cycles, it is advisable to add Hoechst 33342 to each FITC stain. This ensures stable nucleus contrast and minimizes the risk of deterioration in segmentation results during the course of the experiment. FITC staining provides high-contrast and clearly interpretable results across all cycles. At the beginning, a certain amount of autofluorescence of the cells can be detected, but this rapidly decreases with increasing number of bleaching cycles. As a result, almost only the specific FITC signal of the antibody used remains in the further course. Although the absolute signal intensity decreases over the course of the cycles, the signal-to-background ratio increases, improving the overall contrast. (Supplemental Figure **??**) The signal remaining after the bleaching process was reliably eliminated during post-processing so that the quantitative analysis of the staining was not impaired. The order of the antibodies used was randomized except for the antibodies that characterize the leukocyte subpopulation (CD3, CD4, CD8, CD16b, CD19, CD33, CD45, CD56, CD66b, CD123, and CD163). The stainability of the cells is maintained across all cycles. The results of the registration confirm the reliability and accuracy of the registration method used for evaluating the experimental image data. The segmentation of the cell nuclei proves to be extremely stable, which is largely due to the consistently high quality of the nuclear stain. This enables reliable and consistent identification of the cell nuclei. The performance of the segmentation algorithm is particularly evident in the fact that precise differentiation and assignment of cell nuclei is guaranteed in both areas with low and high cell density. With regard to the harmonization of cell identities over several cycles, it can be stated that a high degree of stability and reproducibility is achieved. This result underscores the effectiveness of the segmentation and harmonization methods used to track individual cells over the entire experimental period. The post-processing pipeline uses simple but robust algorithms to provide reliable information about the fluorescence intensities of the respective antibody stains at the single-cell level, allowing this data to be efficiently summarized and evaluated. Since the focus during the development of the algorithms was on precision rather than computing power, there is still room for optimization. Adapting and accelerating the algorithm could save considerable time and open up possibilities for real-time analysis. The individual signal intensities of the different stains cannot be directly compared with each other, nor can the signal intensities of the same antibody across different data sets. For this reason, it is currently only possible to interpret whether the signal of a cell for antibody A is relatively high or low compared to antibody B within the same experiment. Absolute statements about the expression strength of a specific marker or comparisons between different experiments are not meaningful. Against this background, a qualitative evaluation that takes into account the relationships between antibody signal intensities appear to be the most sensible approach. By comparing the relative signal intensities within a data set, patterns in the expression landscape can be revealed and functional and phenotypic differences between cell groups can be identified. The analysis of the expression patterns of different surface markers in the SPADE tree clearly illustrates the phenotypic heterogeneity of the cell population under investigation. The graph-based representation allows clusters with similar expression profiles to be identified and their relationships to each other to be analyzed. In this way, subpopulations with characteristic marker expressions can be identified and their potential functional relevance in the context of the biological system under investigation can be deduced. In the long term, it would be desirable to improve the quantitative comparability of signal intensities both within and between datasets through appropriate normalization strategies or reference standards. Until then, the qualitative analysis of expression patterns provides a valuable basis for characterizing cellular diversity and evaluating the method itself. Whether the observed antibody stainings actually reflect biological reality can only be conclusively assessed by future experiments. Cross-validation via fluorescence-activated cell sorting (FACS) analyses and the inclusion of a suitable control group are planned for this purpose. Until then, the Ultra-Content Screening method provides a new approach for analyzing blood samples and obtaining more information about leukocyte differentiation and distribution. It has the potential to enable comprehensive molecular characterization in the future, a prospect that could prove instrumental in facilitating diagnostics and therapy planning.

## Acknowledgements

We would like to thank Gabriel Crespo López-Urrutia for his preliminary work in developing the method.

## Availability of data and materials

The datasets analyzed during the study are available from the corresponding author on reasonable request.

