## Supplementary Figures for "Ultra-Content Screening (UCS): Toward the Big Blood Picture"

### Supplements

Supplemental Table 1: Patient information included in the results. The data include sex, AML karyotype, hotspot mutations, ELN classification, cell count before freezing, cell count after fixation and related yield percentage. Complex karyotype of patient sample 3: 45,XX,-7[15]/45,idem,t(2;4)(p23;q25)[2]/45,idem,del(4)(?q31),+?14,-21 (EVI-translocation).

| Sample List | | | | | | | |
|---|---|---|---|---|---|---|---|
| Patient Sample | Gender | Karyotype | Hotspot Mutations | ELN-2022 | Initial cell count [mio.] | Fixation Yield [mio.] | Yield Percentage |
| 1 | Male | 46, XY | DNMT3A, FLT3, NPM1 | 2 | 75 | 28.4 | 37.87% |
| 2 | Male | 46,XY,del5q(q31q35) | NA | 3 | 25 | 17.925 | 71.70% |
| 3 | Male | Complex Karyotype | DNMT3A | 3 | 100 | 29.96 | 29.96% |
| 4 | Male | 46,XY,t(8;21) | - | 1 | 30 | 27.27 | 90.90% |
| 5 | Male | 46, XY | NRAS, CEBPA (biallelic), FLT3-ITD (0.1 ratio) | 2 | 69 | 21.42 | 31.04% |
| 6 | Female | 46,XX | CEBPA (biallel), RUNX1, DNMT3A, ASXL1, IDH1 | 1 | 35 | 33.87 | 96.77% |
| 7 | Female | 46,XX | NPM1, FLT3-ITD | 2 | 80 | 62.75 | 78.48% |
| 8 | Female | NA | NRAS, TET2, NPM1 | 1 | 46 | 30.195 | 65.64% |
| 9 | Female | 46,XX | NPM1, FLT3-ITD, FLT3-TKD, WT1 | 2 | NA | 8.4 | - % |
| 10 | Male | 45,X,-Y,inv(10)(p11.2q21) | NPM1, FLT3-ITD, CEBPA, GATA2, TET2 | 2 | NA | 30.20 | - % |
| 11 | Male | 46,XY | ASXL1, EZH2 | 3 | 40 | 31.5 | 78.75% |

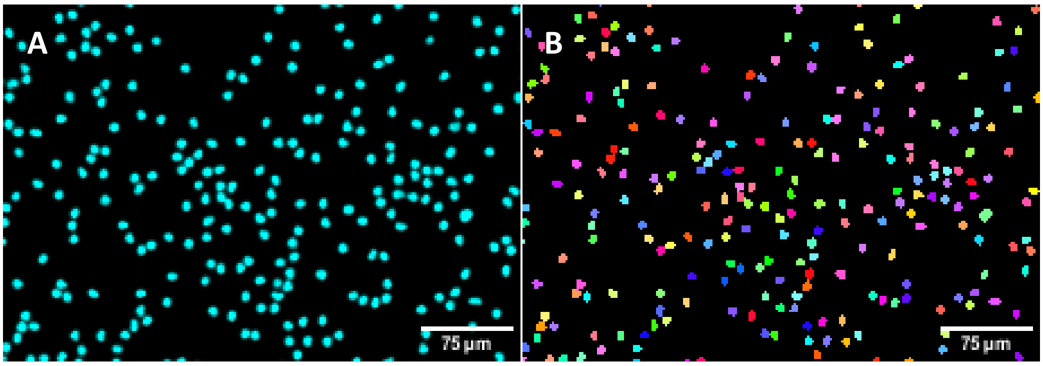


Supplemental Figure 1: Demonstration of Voronoi-Otsu-Labeling segmentation algorithm. (a) Section of a Hoechst 33342 core stain image. (b) Corresponding segmentation result to image section (a).

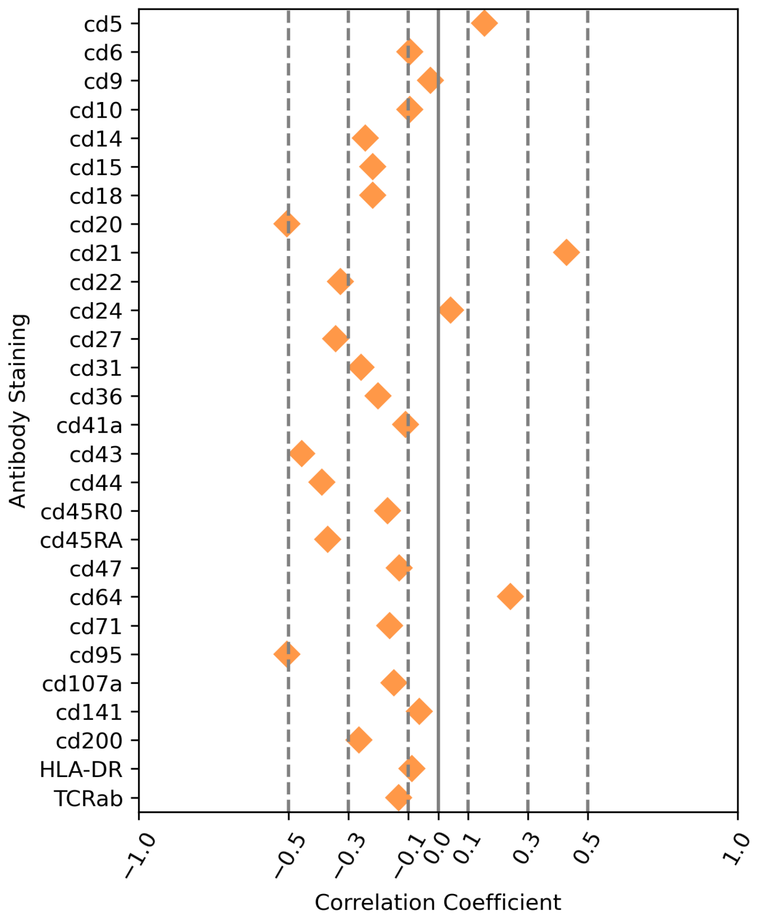


Supplemental Figure 2: Spearman correlation of measured signal intensity in comparison with the position of the staining cycle in the experiment. A negative correlation coefficient suggests that the later the staining is performed in the experiment, the lower the measured signal. Coefficient values of 0 to 0.1: no effect, 0.1 to 0.3: weak correlation, 0.3 to 0.5 moderate correlation, 0.5 to 1: strong correlation. Antibody staining characterizing the leukocyte subpopulations (CD3, CD4, CD8, CD16b, CD19, CD33, CD45, CD56, CD66b, CD123, and CD163) was performed at the beginning of the experiments in the same order. Therefore, these data cannot be taken into account in the correlation.
